# Behavioral assessment of Zwicker tone percepts in rodents

**DOI:** 10.1101/2022.12.22.521554

**Authors:** Achim Schilling, Konstantin Tziridis, Holger Schulze, Patrick Krauss

**Affiliations:** Experimental Otolaryngology, ENT-Hospital, Head and Neck Surgery, University Hospital Erlangen, Friedrich-Alexander University Erlangen-Nürnberg (FAU), Germany

**Author notes:** Both authors contributed equally (Achim Schilling, Konstantin Tziridis). **<u>Corresponding author:</u>** Dr. Patrick Krauss, Experimental Otolaryngology, ENT-Hospital, Head and Neck Surgery, University Hospital Erlangen, Waldstrasse 1, 91054 Erlangen, Germany.

**Keywords:** GPIAS, acoustic startle reflex, aversive conditioning, shuttle box, tinnitus, Zwicker tone, auditory phantom perception

## Abstract

The Zwicker tone illusion can serve as an interesting model for acute tinnitus, an auditory phantom percept still not fully understood. Recent mechanistic models suggest that the underlying neural mechanisms of both percepts are similar. However, to date it is not clear if animals do perceive the Zwicker tone at all, as up to now no behavioral paradigms are available to objectively assess the presence of this phantom percept.

Here we introduce, for the first time, a modified version of the gap pre-pulse inhibition of the acoustic startle reflex (GPIAS) paradigm - usually used to assess the presence of a tinnitus percept in animals - to test if it is possible to induce a Zwicker tone percept in our rodent model, the Mongolian gerbil. Furthermore, we developed a new aversive conditioning shuttle box learning paradigm and compare the two approaches.

We found a significant increase in the GPIAS effect when presenting a notched noise compared to flat white noise gap pre-pulse inhibition, indicating that the animals actually perceived a Zwicker tone. However, in the aversive conditioning learning paradigm, no clear effect could be observed in the discrimination performance of the tested animals. When investigating the CR+ responses, an effect of a possible Zwicker tone percept can be seen, i.e. animals show identical behavior as if a pure tone was presented, but the paradigm needs to be further improved.

## Introduction

The Zwicker tone illusion was first described as a so called ‘‘auditory afterimage” by Eberhard Zwicker in 1964 (Zwicker, 1964). It is typically evoked by the presentation of white noise with a spectral gap (notched noise, NN) but can also be triggered by e.g. loud low-pass filtered noise (Franosch *et al*., 2003). Although the name auditory after-image indicates some analogies of Zwicker tone and visual phantom percepts, the underlying mechanisms are assumed to be completely different, i.e., not caused by a bleaching of the sensing pigment or by some cortical adaptation (Shimojo *et al*., 2001). Instead, a neuronal mechanism most probably in the early stages of the auditory pathway is supposed to be the cause (Norena *et al*., 1999). In recent years, the Zwicker tone became a model system for tinnitus research as it can be evoked fast and reliably and does not cause harm to the auditory system (Leske *et al*., 2014). Furthermore, as the Zwicker tone is perceived as a pure tone (Fastl & Stoll, 1979) with a pitch within the spectral notch, its frequency can be tuned manually by simply shifting the spectral notch of the presented noise. Within this approach, Parra and Pearlmutter have already shown a correlation of the presence of tinnitus and the probability of perceiving a Zwicker tone after noise presentation, indicating that the underlying neural mechanisms might be similar (Parra & Pearlmutter, 2007). Thus, they describe the Zwicker tone as well as tinnitus as an effect of increased neuronal gain along the auditory pathway. There are several mechanistic explanations for increased neuronal gain based on molecular or cell biology (Norena, 2011; Turrigiano, 2012; Tighilet *et al*., 2016; Gainey & Feldman, 2017). However, all these models have in common that they suppose mechanisms that need hours or days to take effect whereas the Zwicker tone percept emerges within seconds or even faster.

In contrast to the models mentioned above, the recent model by Krauss and colleagues is the only one that proposes a neuronal network based effect, and thereby is capable to operate on very short time scales of seconds or even below. It therefore is the only model that may explain both, the emergence of Zwicker tone and acute tinnitus (Krauss *et al*., 2016; Schilling *et al*., 2021; Schilling *et al*., 2022b). The model predicts a threshold improvement during the perception of a Zwicker tone and/or tinnitus by a physiological mechanism optimizing information transmission from the cochlea into the subsequent auditory pathway. The hearing threshold improvement has already been shown during Zwicker tone perception (Wiegrebe *et al*., 1996), tinnitus (Krauss *et al*., 2016; Gollnast *et al*., 2017), in a new animal model paradigm of long-term notched noise exposure to simulate hearing loss (Krauss & Tziridis, 2021), and in a computational model of the auditory pathway based on deep neural networks (Schilling *et al*., 2022a). Taken together, all these findings indicate that the Zwicker tone paradigm is a valuable model to investigate and understand the tinnitus phantom percept. However, in order to investigate Zwicker tone perceptions in animal models (such as rodents) a behavioral paradigm for objective assessment of the presence of this phantom percept is crucial. Furthermore, this behavioral correlate of Zwicker tone is a basic requirement for the interpretation of neurophysiological data seeking to describe the neuronal mechanisms leading to auditory phantom percepts. The development of such a behavioral paradigm is particularly challenging, as the Zwicker tone percept is transient and fades within just a few seconds after induction.

Here, we introduce and test two potential behavioral paradigms to assess Zwicker tone percepts in rodents. The first one is based on the gap pre-pulse inhibition of the acoustic startle reflex (GPIAS) paradigm, originally introduced by Turner and coworkers in 2006 (Turner *et al*., 2006) to assess possible tinnitus percepts in rodents. There, a gap of silence within a moderate band-pass filtered noise is presented before a startle stimulus (e.g. a white noise burst). The gap of silence, if perceived, leads to a suppression (pre-pulse inhibition, PPI) of the acoustic startle reflex. The idea of the paradigm is that the gap of silence is masked (“filling in” hypothesis) by a tinnitus percept, making the gap less salient and hence decreasing the gap-induced pre-pulse inhibition. Due to observations (for details cf. Discussion) indicating that spectrally different pure tones or phantom percepts may serve as an additionally salient pre-pulse within the GPIAS paradigm (Gaese *et al*., 2009; Schilling *et al*., 2017), we developed the hypothesis that a Zwicker tone induced by a notched noise should also lead to an increased GPIAS. This is because the Zwicker tone is perceived as a pure tone and consequently sounds fundamentally different compared to the notched background noise used, resulting in contrast enhancement. Along these lines, we here propose a behavioral paradigm based on the GPIAS paradigm where the band-pass noise is replaced by broadband, notched noise potentially inducing a Zwicker tone percept. As we assume the Zwicker tone to serve as a salient pre-stimulus, which is perceived in the gap of silence, we consequently predict an increased GPIAS compared to the control condition of a GPIAS induced by a gap of silence in a white noise background. Therefore, the effect should be opposite compared to the masking of the gap-effect expected in tinnitus testing.

In a second, alternative approach, we use an aversive conditioning GO-NOGO paradigm in the shuttle box where animals are trained to discriminate between a white noise stimulus followed by a pure tone (reinforced conditioning stimulus, CS+) and the white noise stimulus followed by silence (non-reinforced conditioning stimulus, CS-). After the conditioning period, the CS+ is replaced by notched noise - presumably inducing a Zwicker tone of the trained frequency - followed by silence. The idea is that in case the animals indeed perceive the Zwicker tone, they should show the same behavior as for the CS+, thereby indicating the perception of a Zwicker tone.

## Materials and Methods

### Animals and Housing

39 male Mongolian gerbils, purchased from Janvier (Le Genest-Saint-Isle, France) were housed in standard animal racks (Bio A.S. Vent Light, Zoonlab, Emmendingen, Germany) in groups of 3 to 4 animals with free access to water and food at a room temperature of 20 to 25°C under a 12h/12h dark/light circle. The use and care of the animals was approved by the state of Bavaria (Regierungspräsidium Mittelfranken, Ansbach, Germany, No. 54-2532.1-02/13).

### Stimulation software

The entire software used to conduct the GPIAS (Gerum et al., 2019) measurements (as well as the evaluation procedures, cf. below) was written in Python 3.6 (Van Rossum & Drake Jr). For basic numerical operations the Numpy library was used (Van Der Walt *et al*., 2011); more complex mathematical operations (e.g., signal filter functions) were implemented using the SciPy package (Oliphant, 2007). Data visualization was realized using the Matplotlib library (Hunter, 2007) and the Pylustrator (Gerum, 2019). Efficient storage of the data was achieved by the usage of the Pandas library (McKinney, 2010).

The software for the shuttle box was implemented in Pascal (Wirth, 1971) and is described in detail in earlier work (Schulze & Scheich, 1999). All data evaluations were performed by hand and transferred to Statistica (StatSoft Europe, Hamburg, Germany); for details see below.

### GPIAS measurements

For the GIPAS measurements on 24 male gerbils a custom-made open-source setup was used as detailed before (Gerum *et al*., 2019). In short, the animals were placed in an acrylic glass restrainer tube, closed with a wire mesh at the front side and a cap at the back end, and placed on a sensor platform fixed to a vibration-proof table. Movements of the sensor platform were registered using a 3D acceleration sensor. Two loudspeakers were placed at a distance of 10 cm in front of the animal. They present, first, the 115 dB SPL startle stimulus (Neo 25 S, SinusLive, noise burst 20ms, flattened with 5 ms sin^2^ ramps) and, second, the 60 dB SPL white / notched noise background (CantonPlus XS.2). Notched (spectral notches centered at 2 kHz or 5 kHz ± half octave; 12 animals each) as well as white noise backgrounds were presented with and without a gap of silence of 50 ms (flanked by 20 ms sin^2^ ramps, 10ms complete silence) stating 100 ms before the startle stimulus.

Three measurement sessions were performed over the course of 14 days for each animal. During the measurements, animals were allowed 15 min of habituation in darkness in the tube. Before the real stimuli were presented, five habituation stimuli were given to “level” the startle responses. Each noise-type stimulus was repeated 30 times with and without gap, summing up to 120 stimuli (**Figure 1**), which took roughly 30 min.

**Figure 1:**
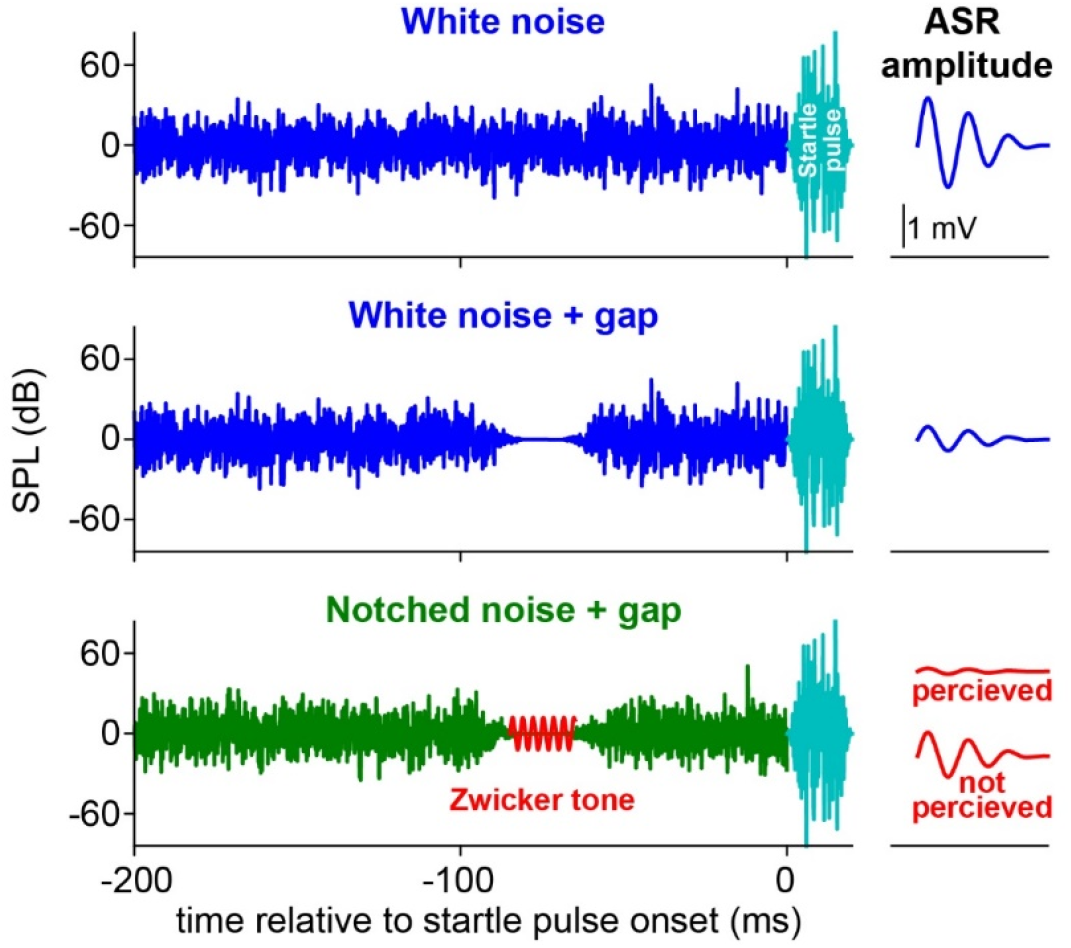
Stimuli for the modified GPIAS paradigm for Zwicker tone detection. The animals were presented with four different stimuli in a pseudorandomized manner. First, a white noise background stimulus without a gap (**top panel**) leading to a large auditory startle response (ASR) after the startle pulse. Second, a white noise background stimulus with a silent gap (**center panel**), leading to a small ASR response due to pre-pulse inhibition (PPI). Third, a notched noise background without gap (not shown), leading to a similar response as in the corresponding white noise condition and, fourth, a notched noise background with gap (**bottom panel**), leading to either a “normal” PPI, when no Zwicker tone is perceived or an increased PPI if the phantom tone is perceived.

The complete evaluation of the GPIAS measurements were performed using custom-made Python programs. The GPIAS effect was quantified by calculating the median from the full combinatorial startle amplitudes as a response to gap and no gap pre-stimulus (for details see Schilling *et al*., 2017). Statistics on the mean GPIAS results were performed with Statistica (cf. below).

### Shuttle box training and testing

The second behavioral paradigm for Zwicker tone assessment was realized as an aversive conditioning GO-NOGO learning paradigm. Another 15 animals were trained in a two compartment shuttle box (both compartments separated by a 6 cm high hurdle) to distinguish between two auditory stimuli (conditioned stimuli, CS) using an electrical foot shock (which must be aversive but never painful) as unconditioned stimulus (US). The animals were supposed to respond to the CS- by staying within their current compartment while during CS+ presentation they were supposed to cross the hurdle into the other compartment (for details see Depner *et al*., 2014). First, the animals were trained for up to 18 days (dependent on the performance of the individual animal: median [interquartile range]: 15 d [9 d, 16 d]). The task to learn was to distinguish between 4 s of silence after 8 s of 60 dB SPL white noise (with 2 ms sin2-ramps; CS-) and 8 s of 60 dB SPL white noise followed by a pure tone (2 kHz, 60 dB SPL, CS+) with a duration of 4 s. The stimuli were created with Audacity, an open source software tool for audio applications. Both stimuli were repeated 30 times each in a pseudorandomized manner in the daily training sessions. The animals were given 2 s to decide if a CS+ tone was presented and cross the hurdle, else a foot shock was given for maximally 2 s. If they crossed the hurdle during the 4 s of CS- silence presentation, a 500 ms foot shock was given. The difference of correct reactions (CR+) and incorrect reactions (CR-) was defined as discrimination performance (DP) which in our case can have values between −30 and +30, where zero indicates no discrimination between both stimuli. After successful completion of the training phase defined as three consecutive days with p values smaller than 0.01 in a chi^2^ test of jump response results, animals were exposed to a single test session. There, the same CS- (white noise with silence) and an adjusted CS+ (notched noise with spectral notch at 2 kHz center frequency, half octave width, followed by silence) were applied according to the same temporal protocol as during training sessions yet but without foot shocks. Our hypothesis was that the notched noise induces a Zwicker tone around 2 kHz which is then perceived as pure-tone stimulus, similar to the one the animals were trained to. Thus, a significant discrimination between CS+ and CS- within the test session would be an indication of a Zwicker tone percept (**Figure 2**).

**Figure 2:**
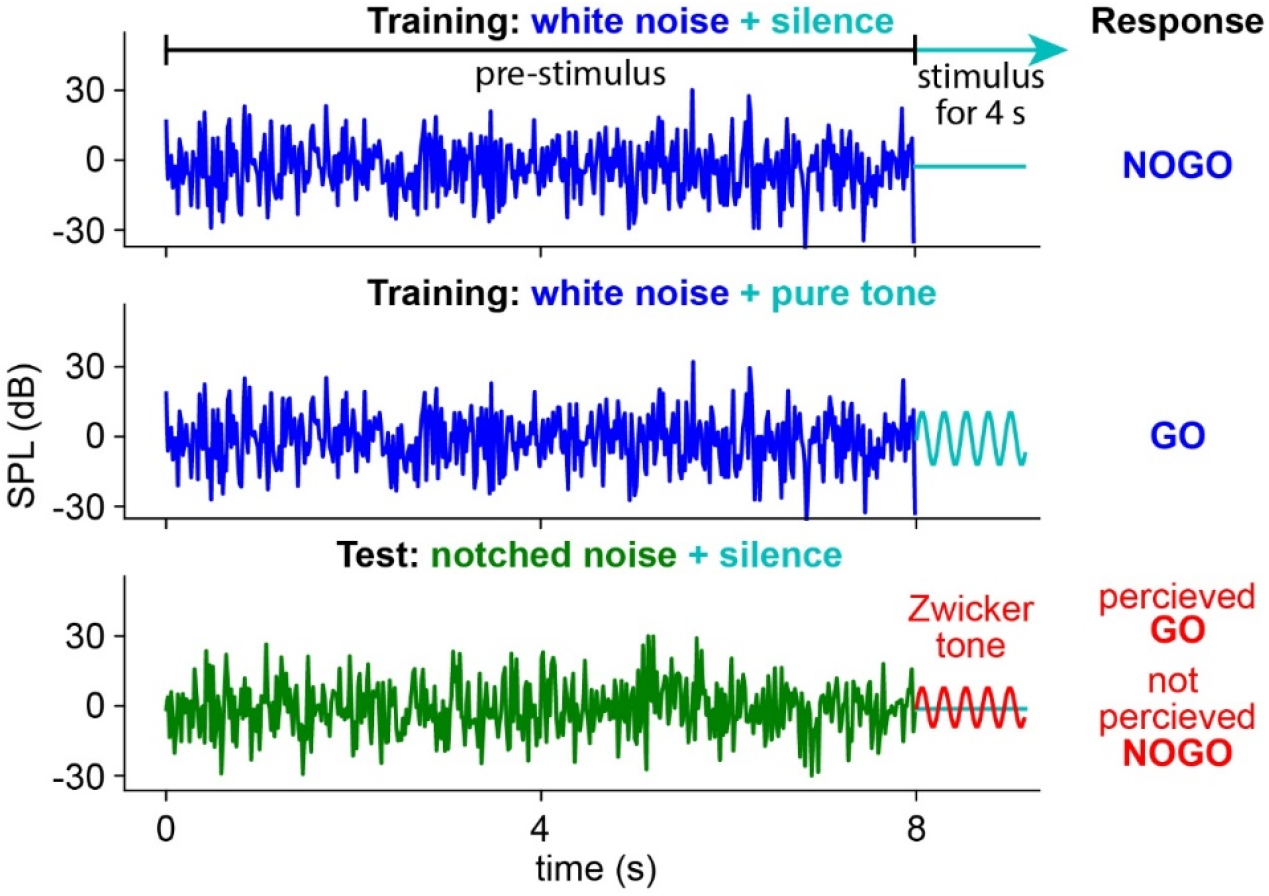
Stimuli of the conditioning paradigm for Zwicker tone detection in the shuttle box. The animals were trained to discriminate between white noise followed by silence (**top panel**) and white noise followed by a pure tone (**center panel**). After successful training, the pure tone was replaced by a potential Zwicker tone by presenting a notched noise with silence (**bottom panel**) instead of white noise with pure tone for one single test session; the white noise with silence stimulus was kept the same. If the animal perceived the Zwicker tone that roughly sounds similar to the GO stimulus, it should respond accordingly.

### Statistics

As mentioned above, all statistics were performed with Statistica. For the evaluation of the GPIAS results, the means of the effect size of the individual gap / no-gap amplitude responses of white noise (WN) and notched noise (NN) were taken for each animal and each measurement and analyzed by Friedman-ANOVAs. Pairwise tests of WN and NN results within one measurement were performed by Wilcoxon matched pair tests. For the comparison of the responses of the animals during the three successful training days and the test day, we analyzed the DP by Wilcoxon matched pair tests and the performance during the first and last ten trials of the sessions separately, as the foot shocks were turned off. This was analyzed by chi^2^ tests.

## Results

### GPIAS paradigm

We exposed two groups of Mongolian gerbils (*12 animals in each group*) to the white noise stimuli with gap, **Figure 1**, top and center panels) and the notched noise stimuli (gap and no gap, Figure 1, bottom panel; notch at 2kHz and 5kHz, respectively, ± half octave) followed by the startle pulse. This experiment was repeated three times (with one week between successive measurements) to check for re-test reliability. The first result we found in this context was the increase in the PPI in the consecutive measurements for both frequency groups (**Figure 3 A and B**). In all four cases (WN and NN in 2 kHz and 5 kHz groups), the significant results of the Friedman-ANOVAs indicate this increase (cf. also **Table 1**). This finding indicates some sensitization of the animals to the gap pre-stimulus. However, the NN-GPIAS compared to WN responses is systematically increased: This is true for both 1^st^ measurements in the two groups (Wilcoxon tests: **Fig. 3A**: p=0.008, **Fig. 3B**: p=0.01) and the 3^rd^ measurement in the 5 kHz group (**Fig. 3B**: p=0.008) while the results in the 2 kHz group were here not different (p=0.16). This was also true for both comparisons in the 2^nd^ measurements (**Fig. 3A**: p=0.34; **Fig. 3B**: p=0.27). This means that, when choosing the right stimulus (in the case of Mongolian gerbils frequencies around 5 kHz), the used paradigm can reliably elicit Zwicker tone percepts and is furthermore suitable for detecting this percept in rodents.

**Figure 3:**
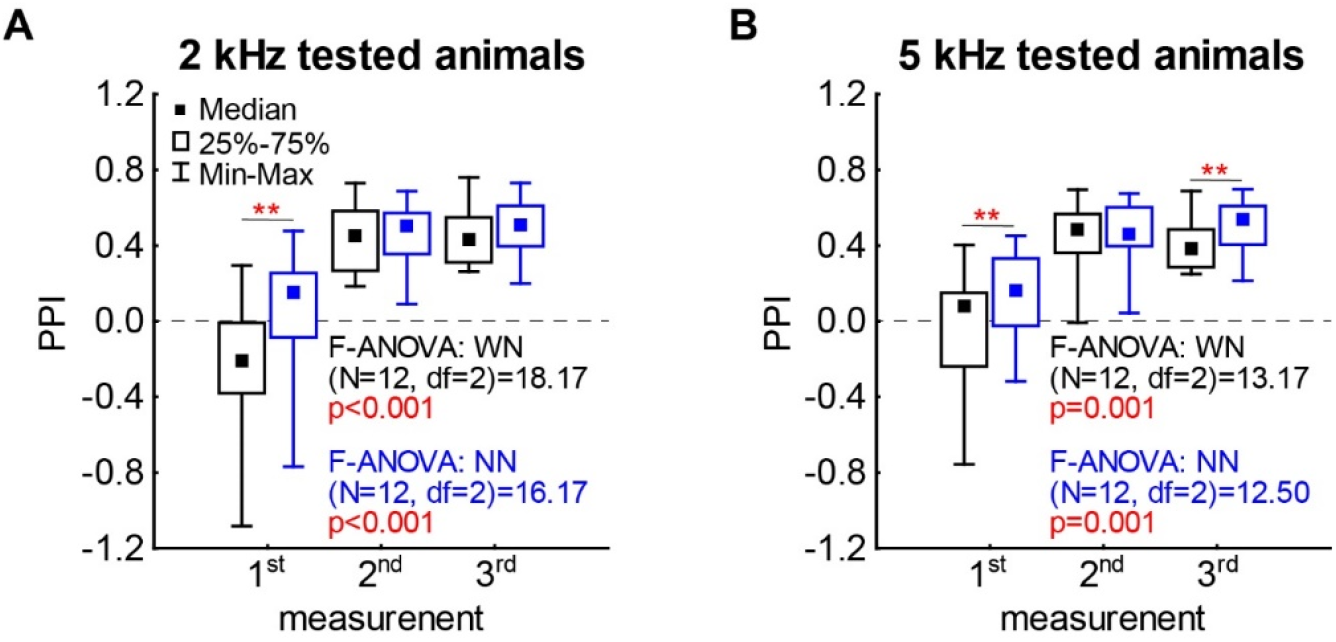
Results of Zwicker GPIAS paradigm. **A** Median of gap induced pre-pulse inhibition (PPI) (full combinatorial, cf. Schilling *et al*., 2017) for the 2 kHz group. Black symbols indicate the values and Friedman-ANOVA statistics for the white noise (WN) condition, blue symbols indicate the results of the notched noise (NN) condition. Asterisks indicate significant differences in the Wilcoxon tests (cf. **Table 1**), ** p<0.01. **B** Median of gap induced GPIAS for the 5 kHz group. Symbols as in **A**.

**Table 1:** Wilcoxon test results for WN – NN comparisons in the three GPIAS measurements

| Frequency | Measurement | Valid N | T score | p value |
| --- | --- | --- | --- | --- |
| 2 kHz | 1 | 12 | 5 | 0.008 |
|  | 2 | 12 | 27 | 0.35 |
|  | 3 | 12 | 21 | 0.16 |
| 5 kHz | 1 | 12 | 6 | 0.01 |
|  | 2 | 12 | 25 | 0.27 |
|  | 3 | 12 | 5 | 0.08 |

### Conditioning paradigm

In conditioning paradigm, we use an alternative approach to elicit and assess Zwicker tone percepts in rodents. In particular, the animals were trained to discriminate between silence and a 2 kHz pure tone after a white noise background stimulus (**Figure 4A**, left panel). After successful training (defined as three consecutive days of a positive DP with p<0.01 in a chi^2^ test), a single test session without any foot shock was performed in order to assess the putative perception of a 2 kHz Zwicker tone. When investigating the DP, the animals did not perceive such a phantom percept, as hardly any animal showed a positive DP with a median of 1 [-3, 3] and a significant performance drop (Wilcoxon test, p<0.001) compared to the three days of successful training DP with 12 [15, 9] (**Figure 4A**, right panel).

**Figure 4:**
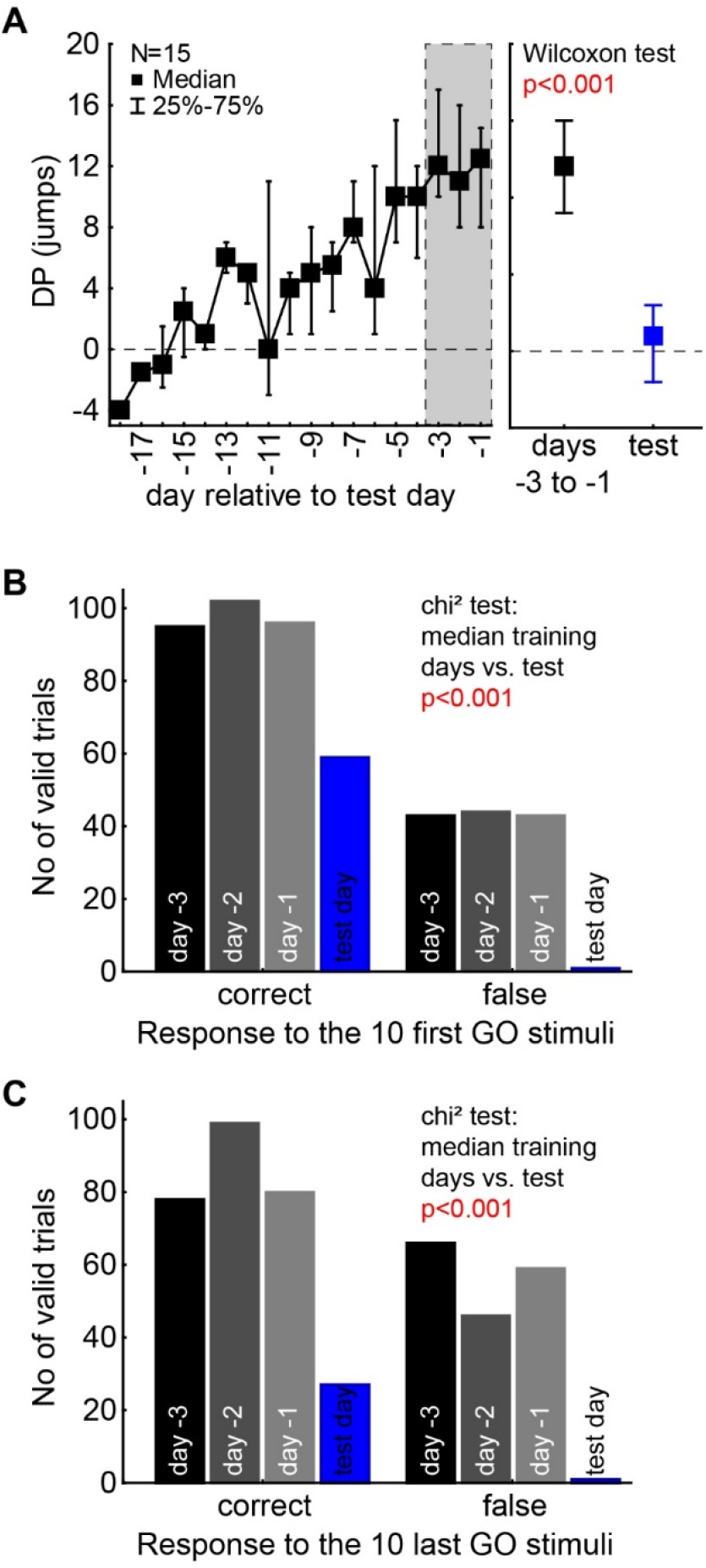
Shuttle box results for Zwicker stimuli. **A** Left panel: Discrimination performance (DP) for training sessions as a function of time relative to test day (day 0). Sufficient learning was defined when the DP was significant on a p<0.01 level in the chi^2^ test over three consecutive days (gray area). Right panel: significant difference (Wilcoxon test) of successful training sessions (black) and test session (blue). **B** Number of correct and false GO-responses for the first 10 GO trials in the successful training sessions and the test session with chi^2^ test result. **C** Number of correct and false GO-responses for the last 10 GO trials in the successful training sessions and the test session with chi^2^ test result.

In order to reveal the reasons for this lack of measurable perception, we inspected the raw data of the total number of correct and false responses to the CR+ in the first 10 (**Figure 4B**) and last 10 trials (**Figure 4C**) in comparison with the responses of the data from the three days of successful learning. In both cases, the chi^2^ tests show differences between the number of responses, with lower correct response rates already in the first 10 trials of the test condition. When comparing the correct responses of the first and last 10 trials separately, we find a significant performance drop in the test condition (Wilcoxon test, p<0.001), while we do not see this drop in the data of the three training days before (Wilcoxon tests, always p>0.05). This indicates that, even though the animals may perceive a Zwicker tone, they do not perform the task, as no foot shocks are applied during the test session.

## Discussion

We here present two novel approaches of behavioral paradigms to objectively assess putative Zwicker tone perception in rodents. The first paradigm is based on GPIAS measurements introduced by Turner and coworkers (Turner *et al*., 2006), while the second is based on an aversive conditioning learning paradigm. In both paradigms, we provide a-proof-of principle that Mongolian gerbils indeed perceive an assessable Zwicker tone illusion, whereas the adjusted GPIAS paradigm yields to the most promising results.

Additionally, the results of that new GPIAS paradigm may have a major impact on the interpretation of the original GPIAS paradigm for tinnitus assessment. This is because the two paradigms differ in the spectral composition of the applied noise: Whereas the original GPIAS paradigm is based on the gap detection ability in band-pass filtered noise, our new Zwicker tone GPIAS paradigm is based on the modulation of the gap detection ability in notched noise. In the original GPIAS paradigm, it is assumed that a potential tinnitus percept leads to a “filling-in” of the gap of silence, and thus to a decreased ability to detect this gap, consequently leading to a decrease of the gap pre-pulse inhibition. However, this mechanism only works, if the perceived tinnitus sounds similar to the surrounding noise.

If the GPIAS paradigm is actually valid for the assessment of a frequency specific tinnitus perception, this paradigm would not predict “filling in” of the gap of silence if the perceived sound is significantly different compared to the surrounding noise. Here, also the difference in presented notch frequencies might play a crucial role (Franosch *et al*., 2003), and may furthermore explain why e.g. the 5 kHz percept is easier to detect compared to a 2 kHz percept. Additionally, it has already been shown that a GPIAS paradigm is at least partially dependent on cortical learning mechanisms (Moreno-Paublete *et al*., 2017; Lanaia *et al*., 2021), which may interact (e.g. suppress) the percept after consecutive presentation. Independent of the frequency or the timing, one can expect an increase of the gap pre-pulse inhibition for the Zwicker tone GPIAS paradigm, as the notched noise should sound completely different compared to the Zwicker tone which resembles a pure tone (Zwicker, 1964), thereby resulting in a contrast enhancement.This is also suggested by experiments presenting an acoustic noise burst or pure tone within a background noise, leading to an increase in the pre-pulse inhibition effect compared to a gap-of-silence approach of equal background sound pressure level (Gaese *et al*., 2009). This increase of the gap detection ability could indeed been shown in our study (cf. **Figure 3**), and thus can be seen as a cross-validation of the original GPIAS paradigm for tinnitus assessment.

On the other hand, the second behavioral paradigm presented in this study, the aversive conditioning learning paradigm in a shuttle box, seemed not suited to assess the perception of a Zwicker tone on the first glance. Only a very small positive DP being not significantly different from zero (sign test, p>0.05) could be found on during the test session. Only when analyzing the responses to the CR+ stimulus alone, an effect for the first 10 CR+ trials of the test day could be found. This indicates a possible Zwicker tone percept, but is clearly only a first hint and proof-of-principle that this approach can be used to assess the existence of the phantom percept. This could be due to several reasons. First, the training may have been too short. We used an individualized approach with defining the training successful when three consecutive days showed a significant DP of p<0.01 in the chi^2^ test. The median number of training days was 15, which is corresponds to the standard time for discrimination training in the shuttle box (e.g., Ohl *et al*., 1999; Ohl *et al*., 2000) but some animals only needed six days to reach this training level, which in retrospective may have been much too early to stop the training. Nevertheless, we did not find a correlation of number of training days and DP on the test day (multiple linear regression analysis, p>0.05). Second, the stimulus may not have been optimal. As we have seen in the Zwicker GPIAS data, the stimulation with 2 kHz stimuli only showed significant results on the first day of testing. The 5 kHz stimulus seems to be much more salient in these kind of paradigms, which we were not aware of when starting the training. Third, even if a strong Zwicker tone was induced by our paradigm, it would only be rated as CD+ by the animals if its pitch was close to the 2 kHz tone used in the training. If on the other hand the pitch of the Zwicker tone would be too different from 2 kHz this would not be generalized into the CS+ category. In a future study, this might be checked by training the animals to generalize any tone pitch to the CS+ category. Fourth and last, as described above we did not use a foot shock reinforcement of the behavior during test day. As we have seen in the raw data of the CR+ response, the animals did respond to the stimuli in the first few trials but soon found out, that no punishment was given when behaving “wrong”, an effect for which the term “extinction learning” has been coined. This and the above mentioned too early end of training may result in the lack of DP on the test day. Nevertheless, a more thorough approach with this paradigm may result in a comparable positive result as the Zwicker GPIAS approach.

Taken together, we were able to show two proof-of-concept approaches to validate a Zwicker tone percept in rodents. With either the easy to use Zwicker GPIAS paradigm or the more elaborate shuttle box paradigm the percept can now be quantified and investigated further.

## Acknowledgements

We are grateful for technical assistance provided by Leonie Stahl, Eva Reingruber, Lea Geissler, Nico Sitzmann and Jana Haag.

## Funding

This work was funded by the Deutsche Forschungsgemeinschaft (DFG, German Research Foundation): grant KR5148/2-1 to PK (project number 436456810) and grant SCHI1482/3-1 (project number 451810794) to AS.

## Notes

### Competing Interest Statement

The authors have declared no competing interest.

